# Barcoding biology: Chemotype predicts variation in genotype, physiology, and stress response

**DOI:** 10.64898/2026.03.22.713522

**Authors:** Rita Ibrahim, Mario González-Jiménez, Jonathan RH Booth, David R Sannino, Alistair O Gemmel, Ignacio Fernandes-Guerrero, Panayiotis Hadjipakkos, Beatriz Castejon-Vega, Rachel Zussman, Nathan Woodling, Klaas Wynne, Adam J Dobson

## Abstract

Predicting biological responses to perturbations such as stress, nutrition, or pharmaceuticals could transform healthcare and biosciences. The task is challenging because complex interactions among many factors interactively drive biological variation, and thereby alter responses. However, metabolism integrates these drivers of variation, suggesting that emergent biological states may be reflected in aggregate chemical states. We define such states as chemotypes and test their predictive utility. Using Fourier-transform infrared (FTIR) spectroscopy coupled with machine learning in *Drosophila melanogaster*, we show that chemotypes serve as proxies for biological variation driven by sex, genotype, nutrition and age, and – critically – predict among-population variation in stress response. These findings indicate that chemotypes provide a computable and integrative representation of organismal biology, predicting genotype, phenotype, and response to perturbation.

## Introduction

Predicting how organisms, patients and populations will respond to perturbations (e.g. diet, pharma, environment) is an unmet challenge in biology and medicine (*1*), which could benefit personalised medicine, agriculture and environmental biology. Responses differ because of biological variation, which emerges from nonlinear interactions among many inputs including genetics, environment and nutrition (*2–5*). The complex basis of variation implies that “bottom-up” predictive strategies likely miss key interactions: biological systems are not the sum of their parts.

An alternative strategy is to proceed not from inputs, but to classify outputs, using proxies of emergent biological states. If variation emerges from the integrated activity of underpinning systems, then those systems likely take on definable states that encode their future trajectories, including response to perturbation. The challenge is to identify a measurable representation of such states that is generalisable, robust, and predictive. Cellular life is fundamentally a chemical phenomenon. We hypothesised that chemical state encodes biological state and, therefore, delineates future responses to perturbation, in a way that can be decoded and barcoded computationally. We extend the concept of “chemotype” to define a computable chemical state, which is expected to integrate diverse drivers of biological variation (*6*). This framework circumvents the biological complexity that impedes bottom-up predictive strategies.

To identify chemotypes, we trained machine learning (ML) algorithms with outputs from Attenuated Total Reflection Fourier-transform Infrared Spectroscopy (ATR-FTIR). ATR-FTIR probes the surface of a biological sample (in this study, the cuticle of whole flies), where infrared light excites molecular vibrations characteristic of chemical bonds. The resulting spectra reflect the combined contributions of proteins, lipids, nucleic acids, and polysaccharides, whose relative abundance and conformational states shape band intensity and profile, providing a rapid and inexpensive readout of aggregate molecular composition (*7–12*). Using *Drosophila melanogaster* as a tractable proof-of-principle system, we find that (A) chemotype acts as a proxy for variation driven by sex, genotype, ageing, and nutrition, suggesting a generalizable framework for decoding emergent biological states, and (B) chemotype can barcode populations by resistance to stress, before exposure. The results outline chemotyping as a versatile and deterministic predictor of organismal state, offering a framework for predictive biology and personalised health interventions.

## Results

### Chemotype proxies for sex, genotype, epistasis, nutrition, and ageing

First, we tested whether chemotyping can discriminate sex, which we anticipated would drive robust chemotypic variation. We tested whether ML accurately discriminated sex-specific FTIR chemotypes, using flies from 12 populations (*13*) to discern genotype-nonspecific signatures (Figure 1A). Our ML pipeline automatically selected features (XGBoost), which were used to train classifiers. Cross-validation analyses (k=20) were used to validate models. In all analyses, Support Vector Machines (SVM) consistently outperformed other classifiers (see Supplementary Materials), and so SVM alone is reported.

**Figure 1.**
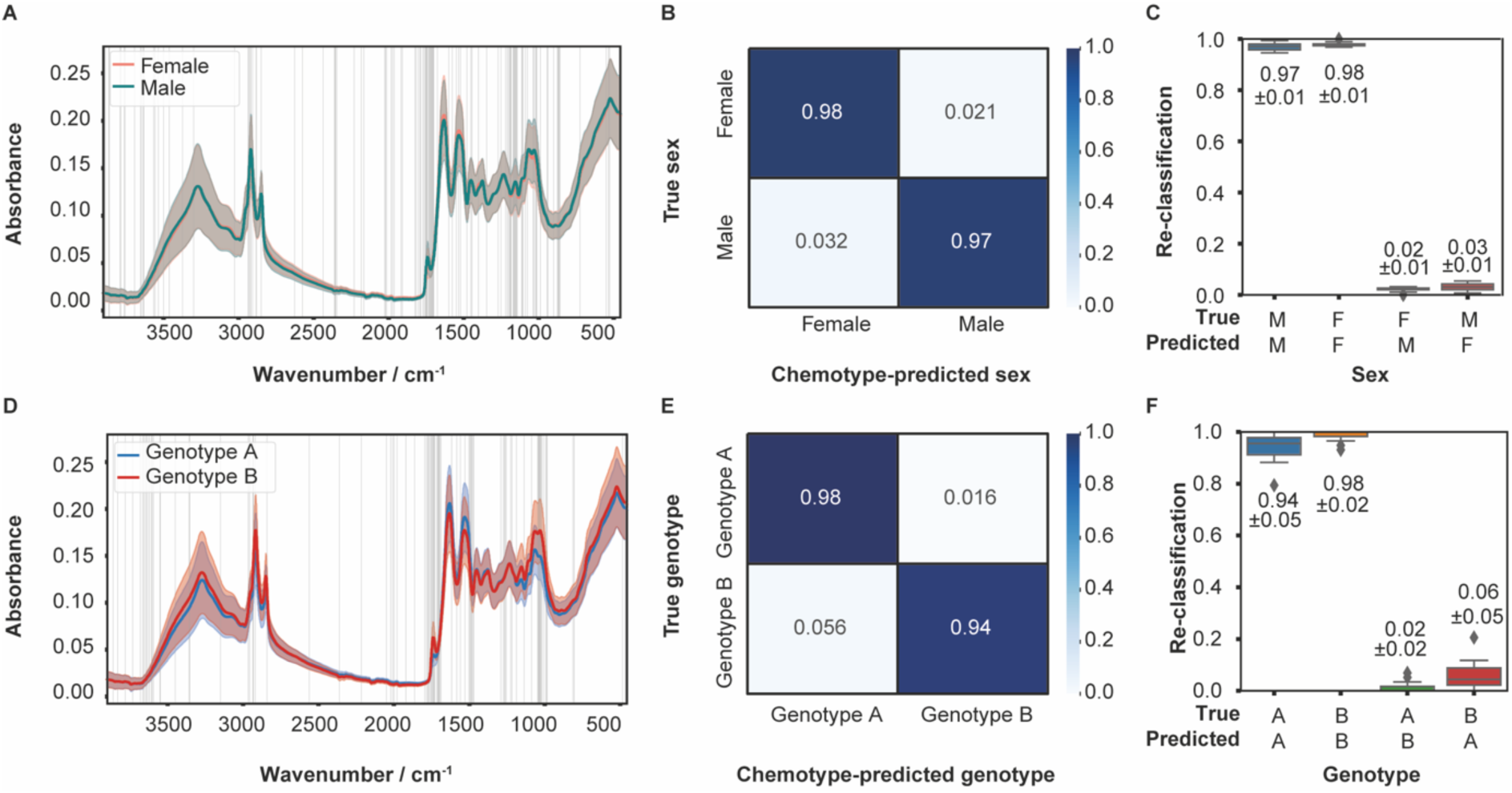
ML classifier learns chemotypes that barcode sex and genotype. (A-C) Male and female flies have machine-learnable chemotypes. (A) Mean FTIR absorbance spectra (±1SD) of Male (N=740) and Female (N=944) flies (pool of twelve populations to ensure genotype-nonspecific chemotype of sex). Vertical lines indicate features selected by XGBoost to discriminate biological conditions. (B) Cross-validation shows that ML classifier learns chemical barcode of biological sex. Confusion matrix shows average proportions males/females correctly/incorrectly reclassified by SVM classifiers of male/female spectra, where models are trained on 80% and validated on 20% of the data. (C) Boxplots showing distribution of SVM reclassification (as per B), for all 20 model iterations. Non-overlapping distributions confirm classifier robustness. (D) FTIR spectra of females from genetically distinct populations originally wild-caught in Australia (genotype *A*, N=288) or Benin (genotype *B*, N=169) were analysed. (E-F) confusion matrix and boxplot generated as per (B-C). Figure S1 evaluates alternative classifier algorithms.

Chemotype barcoded 98% of females and 97% of males (Figure 1B), with negligible errors (Figure 1C). Spectra were not strikingly different by eye, and means were not significantly different at every feature selected for classifier training (Figure S1B), suggesting that ML discriminated the chemotype signature of sex from higher-order patterns. We then tested whether chemotype barcoded genetic differences among populations, which also drive significant biological variation. Chemotype accurately barcoded 98% of flies (females) from an Australian (*A*) population, and 94% of flies from a Beninese (*B*) population (*14*) (Figure 1D-F). Thus, it appears that chemotype can serve as a proxy for both sex and genotype.

We tested the versatility of chemotyping by asking whether we could detect signatures of more complex genetic and physiological sources of variation, specifically genetic interactions (epistasis), sex-specific epistasis, age, diet, and age-by-diet effects. We were able to classify all sources of variation tested (Table 1, Supplementary Text 2-3). Gene–gene interactions (i.e. epistasis between alleles) can generate robust biological variation, with the effect of one variant depending on another. This can be exemplified by interactions between variants of genes on mitochondrial DNA (mtDNA) and variants of genes on nuclear DNA (nDNA). We combined mtDNA and nDNA from the *A/B* populations (Figure 1D-F) (*13*), generating variation that we have shown to alter responses to nutritional and pharmacological interventions (*15*, *16*), and which can potentially be modified further by sex differences (*17*). Chemotypes classified all combinations of sex, mtDNA and nDNA tested, and could also detect mtDNAs and nDNAs (at even higher rates) (Table 1, Supplementary Text 2). To test whether chemotype could also discriminate non-genetic biological variation, we aged flies and changed nutrition (Table 1, Supplementary Text 3). Chemotyping accurately barcoded age, anti-ageing diets (i.e. dietary restriction (*18*, *19*)), age-specific effects of anti-ageing diets, and high-fat diet (Table 1), indicating capacity to discriminate inducible physiological differences, including the emergent properties of interacting factors. Altogether we conclude that chemotyping is a flexible framework that discriminates diverse drivers of biological variation.

**Table 1.** Chemotype discriminates biological variation driven by a wide range of predictive variables. Flies were subjected to each of the given sources of variation (Sex and Genotype as per Figure 1) and chemotyped. Cross-validation was performed to generate a performance index for SVMs trained to chemotypes representing each source of variation, and compared to the null reclassification rate (i.e. expected rate of correct reclassification at random). SVMs were able to accurately discriminate chemotypes representing all sources of variation tested.

| Variable | SVM performance index |  |
| --- | --- | --- |
|  | Observed<br>(mean±1SD, k=20) | Null<br>(1/n. categories) |
| Sex | 0.97±0.01 | 0.5 |
| Genotype (A vs B) | 0.97±0.02 | 0.5 |
| mtDNA (AA vs BA)* | 0.84±0.03 | 0.5 |
| mtDNA (AB vs BB)* | 0.89±0.03 | 0.5 |
| nDNA (BA vs BB)* | 0.98±0.02 | 0.5 |
| nDNA (AA vs AB)* | 0.97±0.01 | 0.5 |
| mtDNA*nDNA | 0.84±0.02 | 0.25 |
| sex*mtDNA*nDNA | 0.79±0.02 | 0.125 |
| Dietary restriction (DR: 5/10/20% yeast) | 0.95±0.01 | 0.333 |
| Age (9/28/42 days) | 0.95±0.01 | 0.333 |
| Age*DR | 0.82±0.02 | 0.166 |
| High fat diet | 0.84±0.06 | 0.5 |

\* Genotypes AA, AB, BA, BB indicate mtDNA and nDNA, respectively, from the founding Australian and Beninese populations. E.g. AB = mtDNA A, nDNA B.

### Chemotype predicts variation in starvation resistance

Having shown that chemotype could discriminate known biological variation, we asked reciprocally whether it could forecast (A) unknown variation (B) in response to a perturbation. We tested how a panel of populations responded to starvation stress. To train a classifier we assayed starvation and FTIR spectra in 108 populations from the *Drosophila* Genetic Reference Panel (DGRP) (*20*, *21*). DGRP flies harbour negligible within-line genetic variation, but substantial among-line variation. Co-housed flies experience the same environment, so allocating co-housed siblings (females) from each line either to chemotyping or starvation stress allowed us to directly associate chemical fingerprints and phenotypic outcome, across the 108 populations (Figure 2A-D). We visualised this association in 2-dimensions with t-distributed stochastic neighbour embedding (tSNE, Figure 2E), which suggested that starvation resistance did indeed correspond to variation in FTIR spectra. Encouraged by this association, we binned populations below/above the 20^th^/80^th^ percentiles as starvation resistant/sensitive (Figure 2F), to isolate their respective chemotypes (Figure 2G). SVMs trained to these chemotypes accurately reclassified 88% of starvation resistant flies, and 84% of starvation sensitive flies, with negligible error (Figure 2H-I).

**Figure 2.**
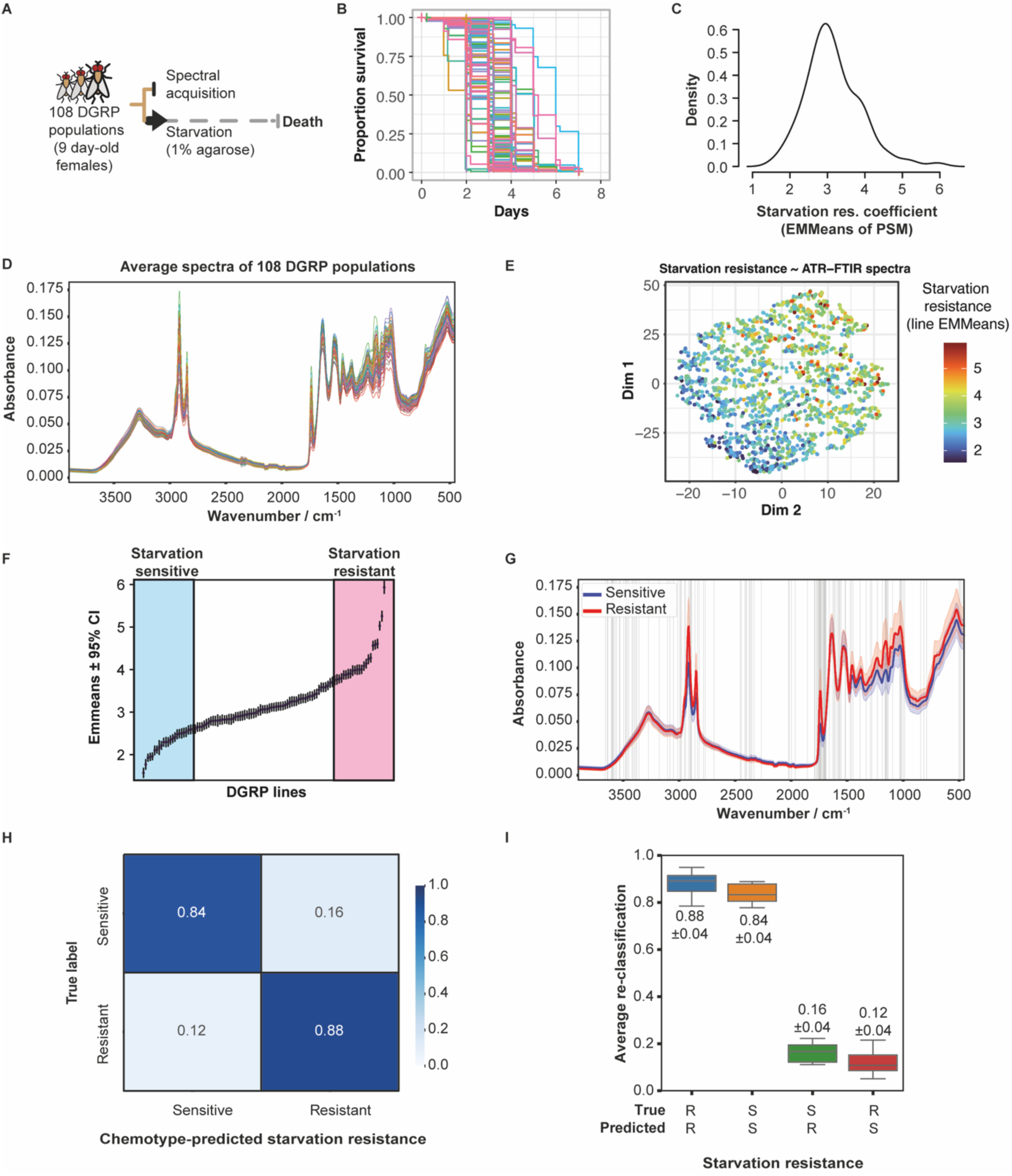
Chemotype is a proxy for variation in starvation resistance. **A.** Experimental design. Females (co-reared siblings) from 108 *Drosophila* Genetic Reference Panel (DGRP) lines were allocated to FTIR spectral acquisition or to starvation, allowing ML to associate genotype-specific spectra and starvation resistance. **B.** Starvation resistance in the DGRP. Plot shows Kaplan-Meier survival estimators for each of the 108 DGRP lines assayed (∼21,000 flies total). Medians (50% survival) are shown with dashed lines. **C.** Variation in DGRP starvation resistance. Variation can be modelled using estimated marginal means extracted from a Parametric Survival Model (PSM). Density plot shows resulting coefficients are variable by genotype (PSM p<0.0001). **D.** FTIR spectra are variable in the DGRP. Each spectrum is the mean of one DGRP line (3,400 flies total). **E.** Variation in starvation resistance corresponds to variation in FTIR spectra. t-distributed stochastic neighbour embedding (tSNE) reduces was used to reduce dimensionality of the spectral data to two axes, with each point representing one individual fly from a given DGRP genotype. Points are coloured by that line’s corresponding starvation resistance coefficient. The correlation across the first dimension indicates that flies with spectra that placed them to the left of the plot are more starvation resistant, and flies with spectra that placed them to the right of the plot are more starvation sensitive. **F.** DGRP line selection for model training. Most and least starvation-resistant DGRP lines are binned according to coefficient of starvation sensitivity. Flies below the 20^th^ quantile (blue) and above the 80^th^ quantile (pink) of the distribution were used. Spectra of these flies were used for model training. **G.** Spectra of flies from the starvation-sensitive and starvation-resistant bins, showing mean (line) and one standard deviation (ribbon). Features selected for classifier training (n=139) are indicated with vertical grey lines. **H.** Cross validation confirms successful model training. The confusion matrix shows the rates at which flies from the starvation-sensitive or starvation-resistant bins were correctly/incorrect reassigned their respective labels. k=20 iterations of cross-validation. **I.** Distribution of correct/incorrect reassignment in model cross validation is shown by boxplots.

Finally, we tested whether chemotypes from DGRP flies predicted starvation resistance *a priori* in independent populations. The independent flies forming the test dataset were from the same populations used in preceding experiments (Table 1, Supplementary Text 2, Figure S3), bearing fully-factorial variation in mtDNA and nDNA, with each of four possible combinations in triplicate (12 populations total) (*13*, *14*, *16*). The classifier predicted two groups among these lines: one starvation-resistant, and one starvation-sensitive (corresponding to nDNA genotype) (Figure 3A). Directly assaying starvation resistance validated this prediction (Figure 3B), with a bimodal distribution of observations (Figure 3C) correlating a bimodal distribution of predictions (Figure 3D). These data indicate that chemotyping can enable *a priori* prediction of responses to intervention, identifying informative signatures that generalize among distinct populations.

**Figure 3.**
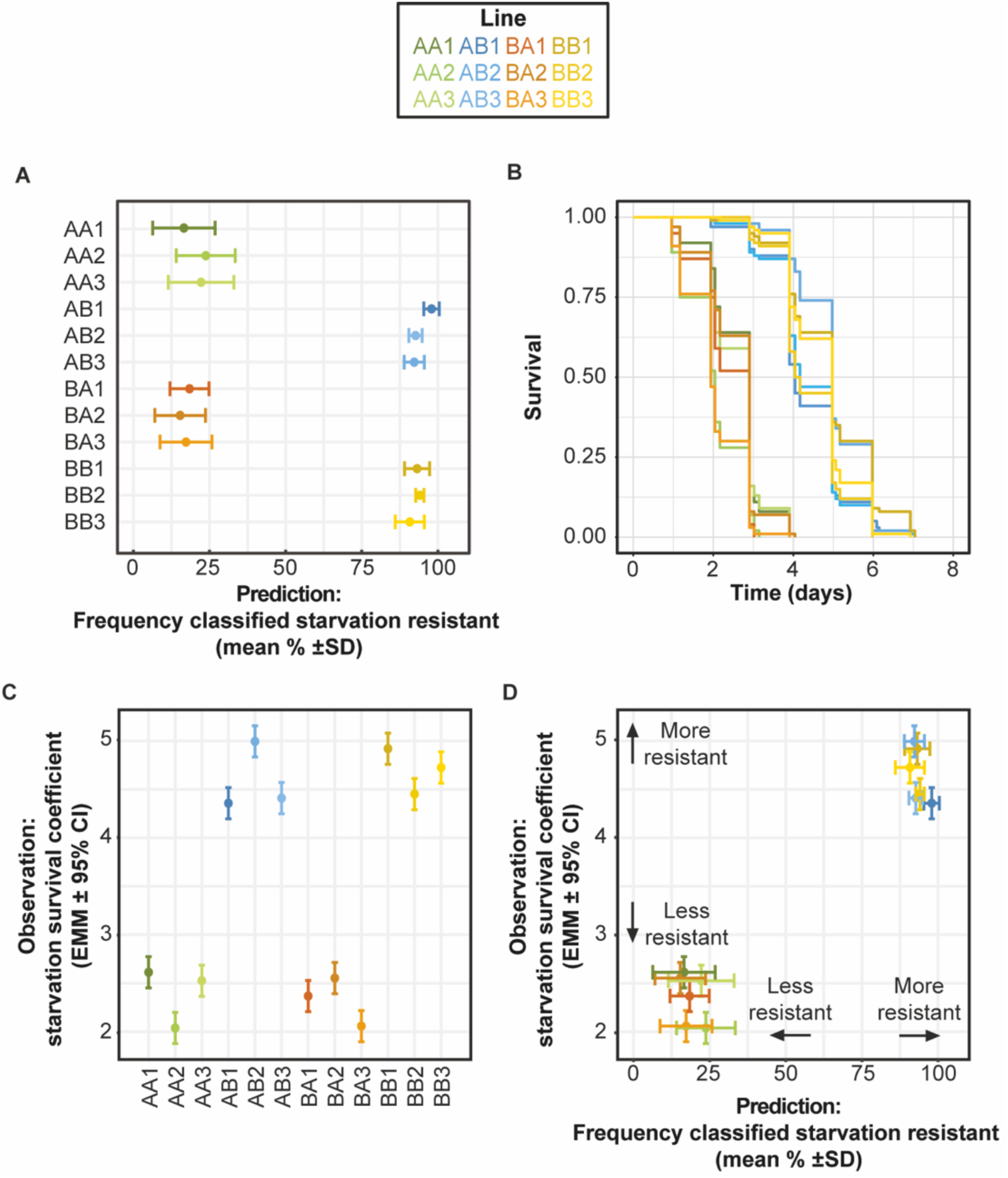
Chemotype predicts starvation resistance. **A.** Classifier trained to chemotypes of starvation resistance (DGRP) makes prediction of starvation resistance in independent lines (mitolines, vis Figure 2). Lines bearing nuclei from Australia (AA1-3, BA1-3) are predicted to be starvation sensitive, while lines bearing nuclei from Benin (AB1-3, BB1-3) are predicted to be starvation resistant. **B.** Testing starvation resistance in mitolines. Plot shows Kaplan-Meier survival estimators for each of the 12 mitolines assayed. Parametric Survival Model p<0.0001. **C.** Estimated marginal means of data shown in B. **D.** Classifier predicts mitoline starvation resistance (Data replotted from A and C). Predicted starvation resistance is the rate at which each of the 12 mitolines were classified as starvation resistant, by the algorithm trained to signature of starvation resistance from DGRP lines. Observed mitoline starvation resistance is given by estimated marginal means of a PSM. The bimodal distribution of classification predicts the bimodal distribution of observation, showing that the classifier was able to accurately predict response to intervention *a priori*.

## Discussion

Taken together, our results demonstrate that chemotypes, captured by ATR-FTIR and decoded by machine learning, contain sufficient information to classify diverse axes of biological variation and, critically, to predict an organism’s response to an upcoming intervention. These findings build on the fundamental concepts that (A) the many diverse drivers of biological variation convergently reconfigure chemistry, and (B) chemical state determines phenotype. We conclude that chemical state can be computed to generate meaningful phenotypic predictions, including variable responses to intervention. Indeed, the presence of biologically-informative chemotypes even in relatively abstracted chemical data, without precisely resolving or quantifying molecular species, suggests that chemical signatures robustly barcode biological complexity. Analytical methods that resolve detailed sample chemistry will likely improve predictive power and provide new insights into the mechanisms of biological variation.

The framework has translational potential. FTIR is rapid, inexpensive, and compatible with minimally invasive human samples e.g. serum, saliva, and urine (*7*, *22*). A longitudinal human study has already shown that chemical/metabolic shifts can precede the clinical onset of metabolic disease by years (*7*, *23*), indicating that chemical signatures may function as early-warning indicators of pathophysiological change. Our new results show that such signatures can also forecast variation in response to intervention. Chemotype-based classifiers could therefore form the basis of individualized medicine, without the risks inherent to using predictors that explain phenotypic variation incompletely (*4*, *24*). By training models to associate chemical signatures with therapeutic outcomes, it may be possible to match patients to the pharmaceuticals, diets, exercise programs, or other interventions most likely to benefit them.

Altogether, our results suggest that the chemical basis of biological variation can be decoded computationally, enabling categorisation of current biological state and, therefore, forecasting of probable future states. We propose this as a generalisable, robust, and scalable framework for predictive biology, which can also be deployed to better characterise mechanisms underpinning biological variation. We foresee implications in healthcare optimization, personalized wellness, and environmental change.

## Data and code availability

All data and code will be made freely and publicly available following manuscript acceptance.

## Acknowledgments

We thank the Medical Research Council (MRC) for support through grants MR/S033939/1, MR/Y019660/1 and MR/P025501/1, the Bill and Melinda Gates Foundation (OPP 1217647), and the University of Glasgow (Lord Kelvin Adam Smith Scholarship and Lord Kelvin Adam Smith Fellowship). This work received funding from the European Research Council (ERC) under the European Union’s Horizon 2020 research and innovation program (grant agreement No. 832703). We thank Matt Piper, Richard Cogdell, Francesco Baldini, Jens Rolff, and members of the Dobson and Woodling labs for constructive comments and discussion; and Alberto Sanz for supporting the project’s practical execution.

## Supplementary materials

**Figure S1.**
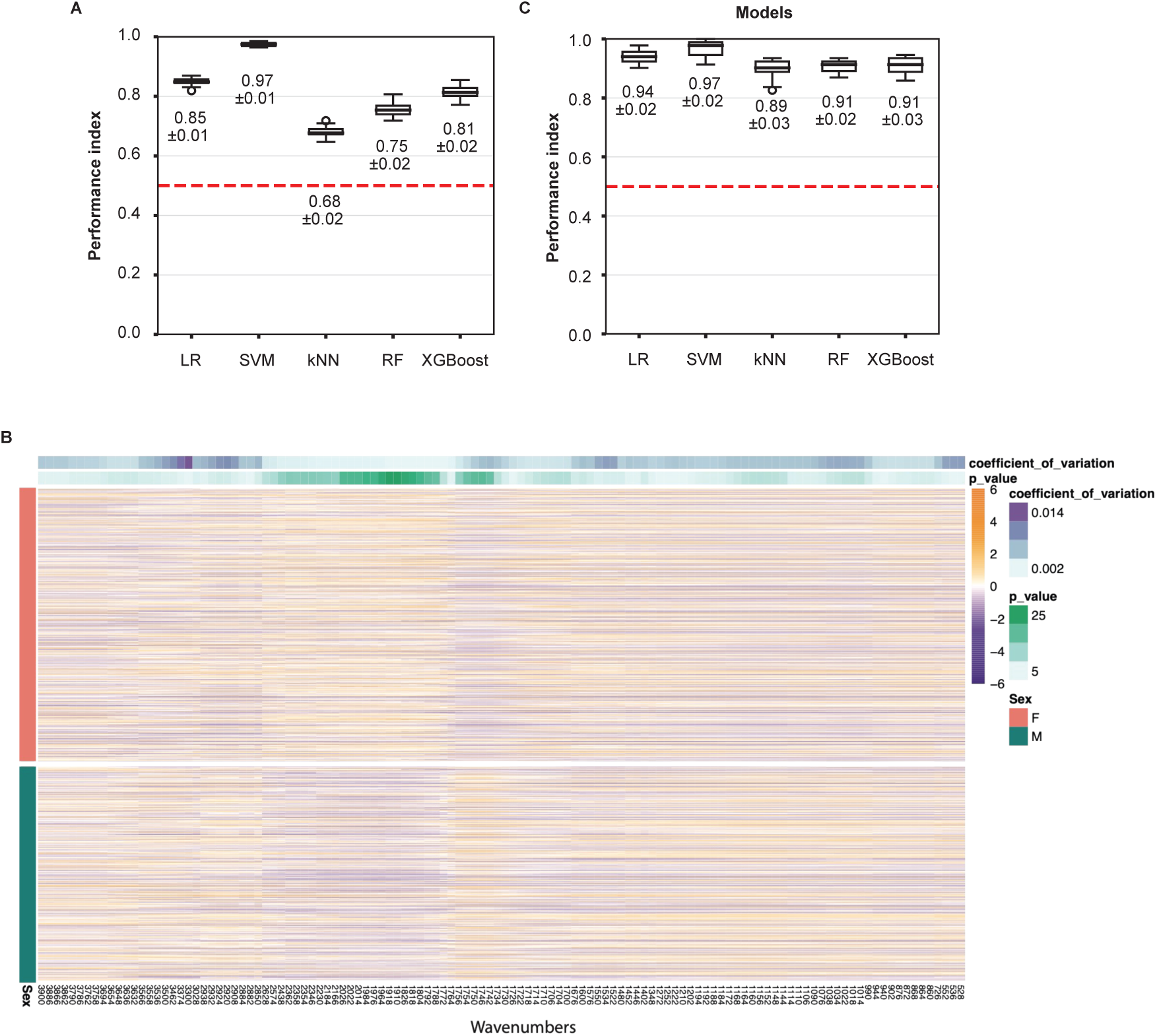
Initial parameterisation of chemotyping in D. melanogaster. (A) Evaluation of five different classifier algorithms for sex. LR=Logistic Regression, SVM=Support Vector Machines, kNN=k Nearest Neighbours, RF=Random Forest. Horizontal red line gives null expected reclassification rate at random (1/2 categories) (B) Heatmap representation of absorbance intensity at features (wavenumbers) selected by XGboost as sex-discriminating. Each row represents one fly, with absorbance values mean-centred and scaled to generate Z-scores. Each column represents a feature. Flies are grouped by sex (left sidebar). For each feature, coefficient of variation and significance of difference (Wilcoxon Rank-Sum p-value) is given at top of figure. Not all features are directionally different between males and females, suggesting ML differentiates the sexes by learning spectral signatures (chemotypes) that are not encoded by mean peak intensities. (C) Evaluation of five different classifier algorithms for genotype.

### Supplementary Text 1 – Machine learning chemotype of sex is non-random

When classifying samples in a dyad – such as female and male – an untrained algorithm is expected to correctly reclassify every second sample at random (i.e. 50% true-positive rate at random). To assess the relative importance of random allocation, we tested how chemotyping reclassified females and males when a third category was available. We took the existing male and female data and generated three subsets, comprising females, males, or a 50/50 mix of males and females which, on average, should present an intermediate chemotype, between male and female (Figure S2A), which we label “DK” (don’t know). We repeated our preceding analysis (Figure 1A) to quantify how often males and females reclassified as DK, and vice-versa. Females and males were reclassified correctly at rates of 0.85 and 0.84, respectively (Figure S2B-C); and both were reclassified as DK at rates of 0.13 (Figure S2B-C). DK flies were reclassified as female or male in approximately equal proportions (0.43 and 0.41, respectively (Figure S2B-C)), which, given that DK were a 50/50 mix of males and females, suggested that the algorithm could assign DK to their original female/male categories. DK, males and females were reclassified as DK at comparable rates (0.13-0.16, Figure S2B-C), suggesting a limit to algorithm performance of approximately 0.145 in the present dataset.

**Figure S2.**
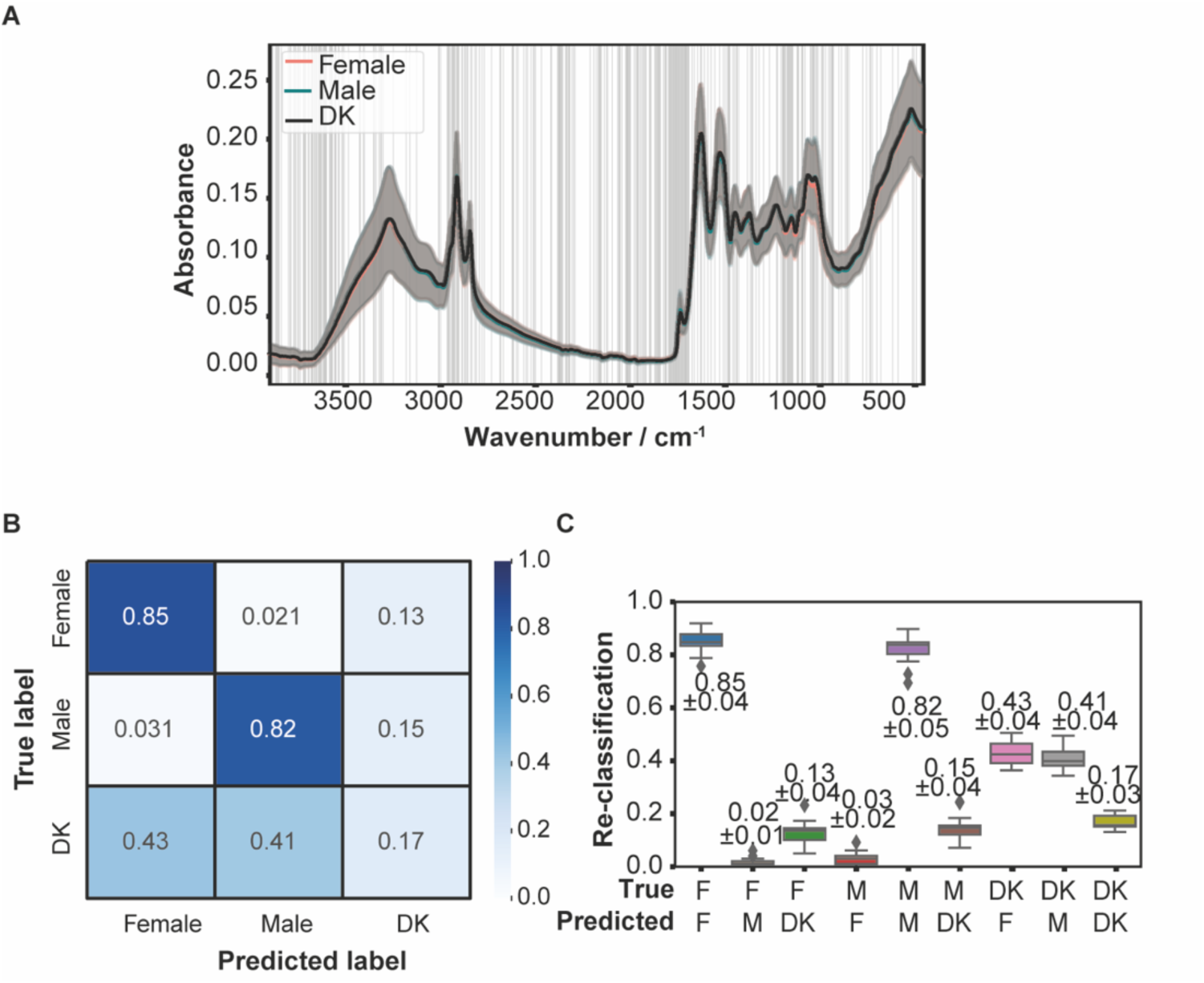
Reclassification of males and females is non-random. (A) Subsets of males and females were binned together to generate a “don’t know” (DK) category. (B) SVM classifiers confronted with the intermediate DK category classify males and females correctly at significantly higher rates than into the DK category; DK flies are reclassified as male/female at approximately equal proportion (∼0.42), reflecting the equal proportions of each biological sex in the DK samples); and flies of all three categories are categorised as DK at approximately equal proportion (∼0.15), giving an apparent limit of classification accuracy for the present dataset. (C) Errors for male/female/DK reclassification are negligible. Boxplots show frequency distribution for correct/incorrect reclassification in k=20 rounds of cross validation. Numbers adjacent to boxes give mean±SD.

### Supplementary Text 2 - Chemotype accurately barcodes complex genetic variation

We examined chemotyping’s capacity to substitute for genetic methods. A key finding in the post-genomic age is that additive effects of genetic variants do not account completely for phenotypic variation (“missing heritability”)(*1*). This suggests that effects of some genetic variants are brokered by interactions with other factors, potentially including other variants (i.e. epistasis). Having shown that chemotyping discriminates chemotypes of geographically-differentiated populations, we asked whether we could detect fingerprints of more subtle and complex genetic variation. Specifically we evaluated signatures of variation on the diminutive mitochondrial genome (mtDNA) and/or more sizeable nuclear genome (nDNA), because epistasis between these two genomes is emerging as a significant source of biological variation (“mito-nuclear” effects), which we have shown to be a significant determinant of response to diet, pharmaceuticals, and altered signalling (*2–4*). We tested whether ML could discriminate signatures of variation in mtDNA, nDNA, and mtDNA*nDNA interactions, using flies from “mitolines” in which mito-nuclear variation has already been characterised(*2*, *4*, *5*), bearing mtDNA and nDNA from Australian (*A*) and Beninese (*B*) populations, in fully-factorial combinations (*AA, AB, BA, BB*, where the first letter denotes mtDNA and second denotes nDNA). Each of the four combinations was replicated independently in triplicate, however for simplicity we here present pooled data from replicates. Furthermore, mito-nuclear effects may also potentially differ between males and females, generating tripartite sex-mito-nuclear variation, so we also evaluated chemical signatures of sex*mtDNA*nDNA interactions.

First we tested whether chemotyping could discriminate chemotype proxies of mtDNA variation, over a fixed nDNA background. The correct mtDNA labels were predicted at rates of 86% (*AA* flies) and 82% (*BA* flies) (Figure S3A-C). Rates were comparable over nDNA *B* (Figure S4). In a reciprocal analysis we tested whether chemotyping could detect fingerprints of nDNA variation, over a fixed mtDNA background, observing accurate prediction at rates of 98% (*BA* flies), and 97% (*BB* flies) (Figure S3D-F), with comparable results for the same nDNAs over mtDNA *A* (Figure S4). These analyses complemented our preceding analysis (Figure 1G-I), suggesting that chemotype can serve as a proxy for genetic variation.

We then tested whether chemotyping could detect chemotypes of mito-nuclear epistasis. Reclassification rates ranged from 80-87% (x̅=84.5%) (Figure S3G-I), indicating accurate prediction of genotype from emergent chemotype, despite its complex, epistatic genetic basis. Finally, we tested capacity to detect sex-mito-nuclear interactions. Reclassification rates ranged from 74-87% (x̅=78.9%) (Figure S3J-L). Altogether, these results indicate that chemotypes can serve as a proxy for genotype, including mitolines in which genotypes are defined by complex genetic interactions.

**Figure S3.**
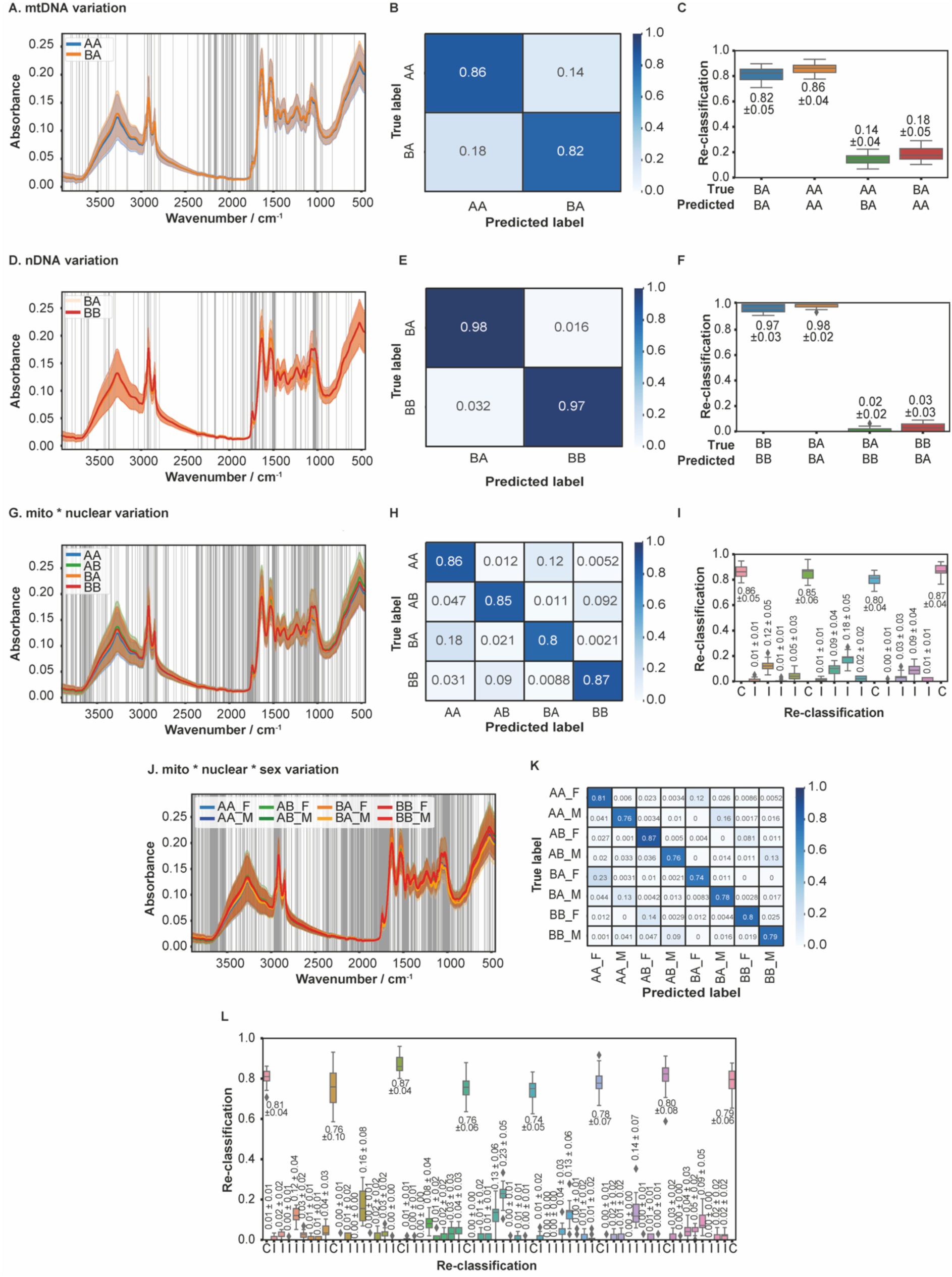
Chemotype accurately barcodes complex genetic variation. Epistatic genetic variation was synthetically generated in flies, by backcrossing nDNA into mtDNA backgrounds. mtDNAs and nDNAs originated from Australia (A) and Benin (B), in fully-factorial combinations (*1*, *2*). (A-C) Genetically distinct mtDNAs have diagnostic chemotypes. (A) FTIR spectra for female flies with genetically distinct mtDNA, over a fixed nDNA background. Mean FTIR absorbance spectra (±1SD) of flies bearing nDNA *A* and either mtDNA *A* (*AA*) (N=288) or mtDNA *B* (*BA*) (N=239). Vertical lines indicate features selected by XGboost to discriminate biological conditions. (B) Cross-validation shows that ML classifier learns chemical barcode of mtDNA variation. Confusion matrix shows average proportions *AA*/*BA* correctly/incorrectly reclassified by SVM classifiers, where models are trained on 80% and validated on 20% of the data. (C) Boxplots showing distribution of SVM reclassification (as per B), for all 20 model iterations. Non-overlapping distributions confirm classifier robustness. (D-F) Genetically distinct nDNAs have diagnostic chemotypes. Plots as per (A-C). (D) FTIR spectra for female flies with genetically distinct nDNA over a fixed mtDNA background. Plot shows mean FTIR absorbance spectra (±1SD) of flies bearing mtDNA *B* and either mtDNA *A* (*BA*) (N=239) or mtDNA *B* (*BB*) (N=169). (E-F) Cross validation shows that ML classifier learns chemotype proxy of nDNA variation. Confusion matrix and boxplot generated as per (B-C). (G-I) Diagnostic chemotypes of mito-nuclear epistasis. (G) FTIR spectra for female flies with genetically distinct mtDNAs and nDNAs, in fully-factorial combinations *AA* (mtDNA *A*, nDNA *A*, N=288), *BA* (mtDNA *B*, nDNA *A*, N=239), *AB* (mtDNA *A*, nDNA *B*, N=248), *BB* (mtDNA *B*, nDNA *B*, N=169). (E-F) Cross validation shows that ML classifier learns chemotype proxy of mtDNA*nDNA variation. Confusion matrix and boxplot generated as per (B-C) and (D-F). (J-L) Chemotype proxies of mito-nuclear-sex interactions. (J) FTIR spectra for female flies with genetically distinct mtDNAs and nDNAs, in fully-factorial combinations. Females are AA_F, AB_F, BA_F, BB_F, males are AA_M (N=145), AB_M (N=165), BA_M (N=183), BB_M (N=247). (K-L) Cross validation shows that ML classifier learns chemotype proxies of mtDNA*nDNA*sex interactions. Confusion matrix and boxplot generated as above. Plots showing feature selection and evaluation of alternative classifier algorithms shown in Figure S3.

**Figure S4.**
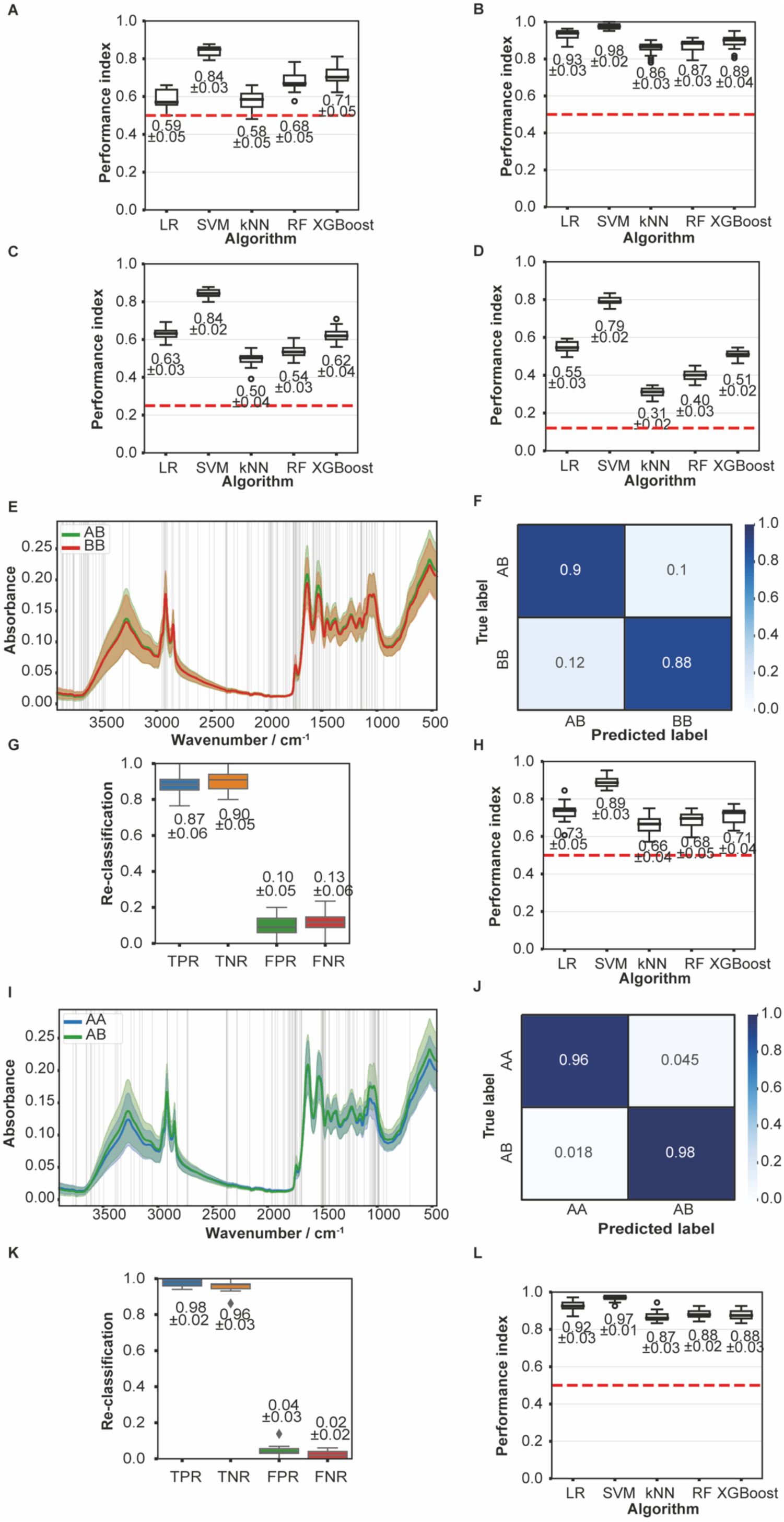
Supporting material demonstrating that chemotype accurately barcodes complex genetic variation. (A-D) Evaluation of alternative classifiers of data presented in Figure S3 A,D,G,J (respectively). LR=Logistic Regression, SVM=Support Vector Machines, kNN=k Nearest Neighbours, RF=Random Forest. (E-L) Confirming chemotype detection of mtDNA variation (E-H, complementing Figure S3 A-C) and nDNA variation (I-L, complementing Figure S3 D-F).

### Supplementary Text 3 - Chemotype proxies of plastic physiological variation

As well as variation in sex and genotype, phenotypic variation also arises plastically in response to intrinsic and extrinsic factors (e.g. age, diet, environment, reproductive status). These factors shape physiology, which in turn shapes response to subsequent interventions. We therefore looked for chemotype proxies for physiological alterations that modulate organismal health.

To generate physiological variation, we manipulated age and diet, since these factors are major causes of variation in health, in humans and other animals. Ageing induces systemic physiological changes, but across animals these effects can be mitigated by moderate dietary restriction (DR)(*6*). DR is thought to reprogram ageing (e.g. *7*, *8*), leading to an expectation that differences between DR and control conditions in youth should be distinct from those in later life, generating age-by-DR effects. We looked for chemotype proxies of DR, age, and age-specific effects of DR.

In flies, DR can be performed by diluting yeast in media(*6*). This form of DR is expected to benefit lifespan, at a cost of reduced fecundity in early life. We confirmed this paradigm in our laboratory finding the expected lifespan and fecundity changes (Figure S5A-B). We therefore tested whether chemotyping could predict these altered physiological states from chemotype proxies. chemotyping was indeed able to discriminate signatures of one week of DR (Figure S5F), at accuracies ranging from 0.93-0.97 (Figure S5G-H). We then tested whether chemotyping could detect signatures of ageing, looking first at diet-independent effects on a fixed diet (10% yeast). Spectra of young (9 days old), middle-aged (4 weeks) and old (6-weeks old) females were visibly distinct (Figure S5C). From these spectra, chemotyping was able to predict accurately how old flies were, with reclassification rates ranging from 0.93-0.99 (Figure S5D-E). Thus, chemotyping can discriminate chemical signatures of age and of diets that modulate ageing.

Next we tested whether chemotyping could identify signatures of DR altering physiological trajectories in ageing flies, i.e. age-by-DR effects. We assayed females after one week of DR (9 days old) and compared flies at the onset of mortality in the high-yeast group (21 days) (Figure S5I). Chemotyping accurately reclassified flies age-by-diet variation at rates ranging from 0.79-0.86 (Figure S5J-K), confirming capacity to discriminate flies that will go on to be longer-lived due to diet-induced plasticity, from those who die young.

In contrast to beneficial dietary changes like DR, some diets can impair health throughout the lifecourse, for example high-energy diets in humans are implicated in rising global rates of metabolic disease. In flies, we have developed a high-fat diet (15% margarine) that reduces egg laying, indicating physiological impairment. We tested whether chemotyping could also detect signatures of this diet-induced malaise (Figure S5L-N), finding that 78% of flies fed high-fat diet were classified correctly, compared to 89% of controls. This result, alongside preceding DR results, indicated that chemotyping can accurately detect chemical signatures of physiological plasticity induced by both beneficial and deleterious dietary change.

**Figure S5.**
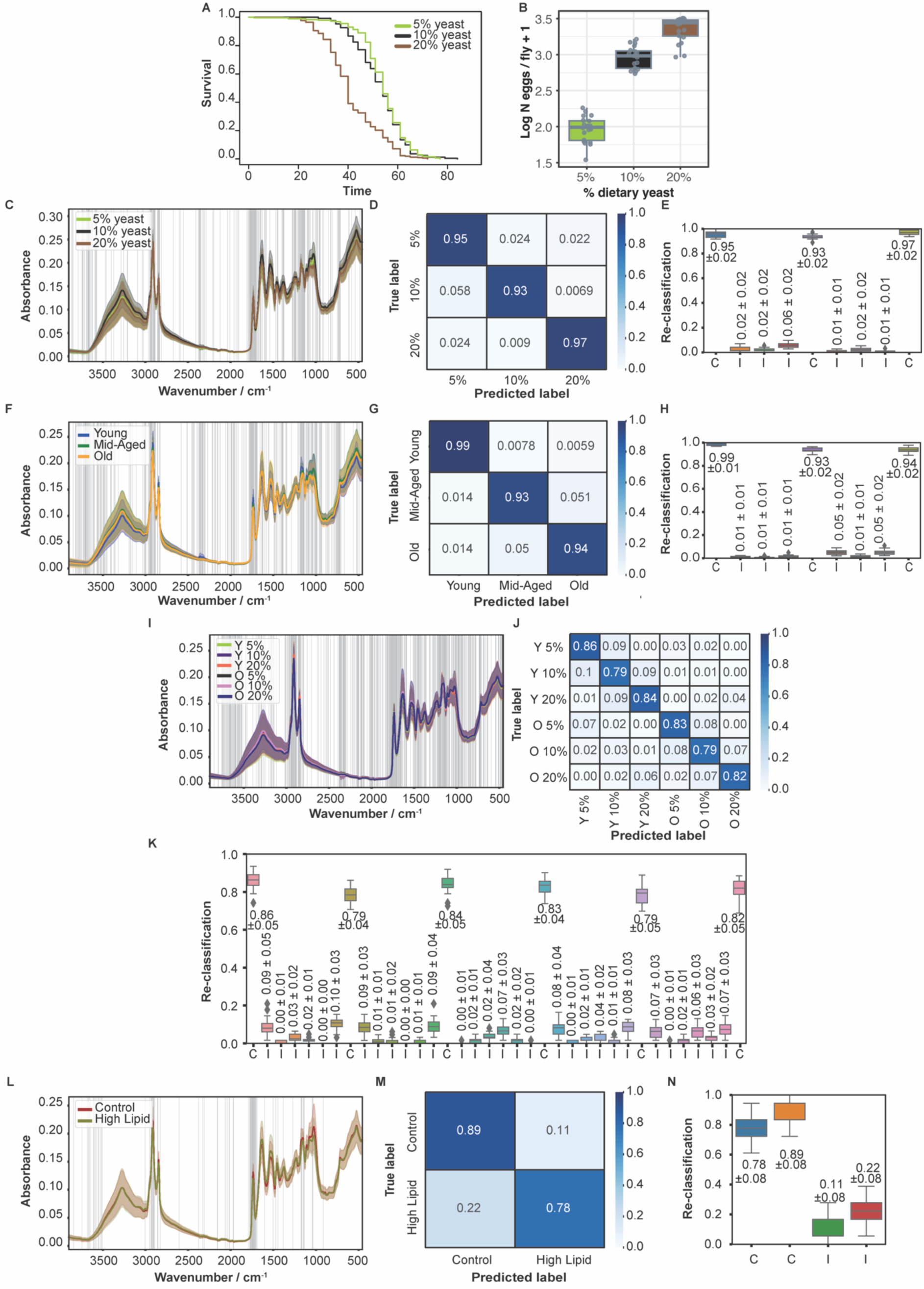
Chemotype accurately barcodes plastic physiological variation. (A-B) Dietary restriction (DR, i.e. yeast restriction) induces anticipated benefits to lifespan (A) and costs to reproduction (B) in female flies. (A) Kaplan-Meier survival plots for flies fed 5% yeast (N=292), 10% yeast (N=257) and 20% yeast (N=312). Parametric survival model (logistic) chi-squared=350.74, p<0.0001. Flies fed 20% yeast were significantly shorter lived than flies fed either 10% or 5% yeast (*Post-hoc* comparisons p<0.0001, 10% vs 5% not significantly different). (B) Boxplots showing eggs laid per vial (∼15 flies per vial; 20 vials per diet) across three yeast levels, after one week of adult-onset DR. Boxplots represent the median, quartiles and 5^th^ and 95^th^ percentiles. Kruskal-Wallis chi-squared=48.53, df=2, p=2.9e-11. All pairwise *post-hoc* comparisons p<0.0001. (C) FTIR spectra for female flies subjected to one week of adult-onset DR (5% N=358, 10% N=358, 20% N=471), showing mean FTIR absorbance spectra (±1SD) per condition. Vertical lines indicate features selected by XGboost to discriminate biological conditions. (D) Cross-validation shows that ML classifier learns chemical barcode of lifespan-altering DR. Confusion matrix shows average proportions of flies from each dietary condition correctly/incorrectly reclassified by SVM classifiers, where models are trained on 80% and validated on 20% of the data. (E) Boxplots showing distribution of SVM reclassification (as per B), for all 20 model iterations. Non-overlapping distributions confirm classifier robustness. (F-H) Differently aged flies have diagnostic chemotypes. Plots as per (C-E). (F) Mean FTIR absorbance spectra (±1SD) of female flies, fed 10% yeast diet, when young (nine days, N=637), mid-aged (four weeks, N=726), or old (six weeks, N=683). (G-H) Cross validation shows that ML classifier learns chemical fingerprint of age. Confusion matrix and boxplot generated as per (D-E). (I-K) Age-specific chemotype for DR. (I) Mean FTIR absorbance spectra (±1SD) of young (Y, 9 days old) and old (O, 21 days old) females fed on 5% yeast (Y 5% N=311, O 5% N=306), 10% yeast (Y 10% N=323, O 10% N=312), and 20% yeast (Y 20% N=311, O 20% N=307). (J-K) Cross validation shows that ML classifier learns chemical fingerprint of age*diet interaction. Confusion matrix and boxplot generated as per (D-E). (L-N) Chemotypes of high-lipid diet (HLD, 15% v/v margarine). (L) Mean FTIR absorbance spectra (±1SD) of control (N=88) and HLD (N=89) females after one week of adult-onset feeding. (M-N) Cross validation shows that ML classifier learns chemical fingerprint of dietary lipid. Confusion matrix and boxplot generated as above. Plots showing feature selection and evaluation of alternative classifier algorithms shown in Figure S5.

**Figure S6.**
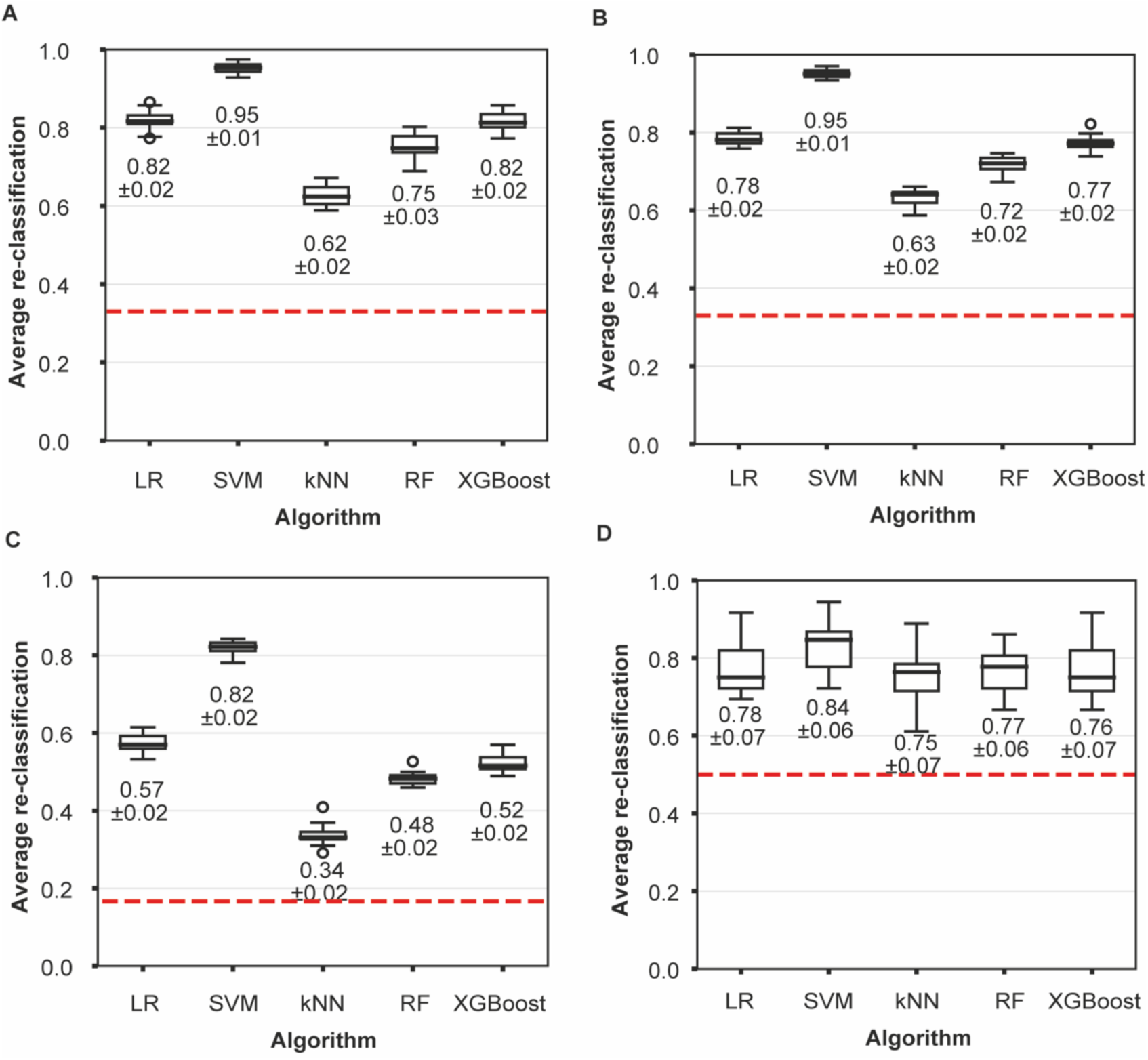
Evaluation of alternative classifiers of data presented in Figure S5 C,F,I,L (respectively). LR=Logistic Regression, SVM=Support Vector Machines, kNN=k Nearest Neighbours, RF=Random Forest.

**Figure S7.**
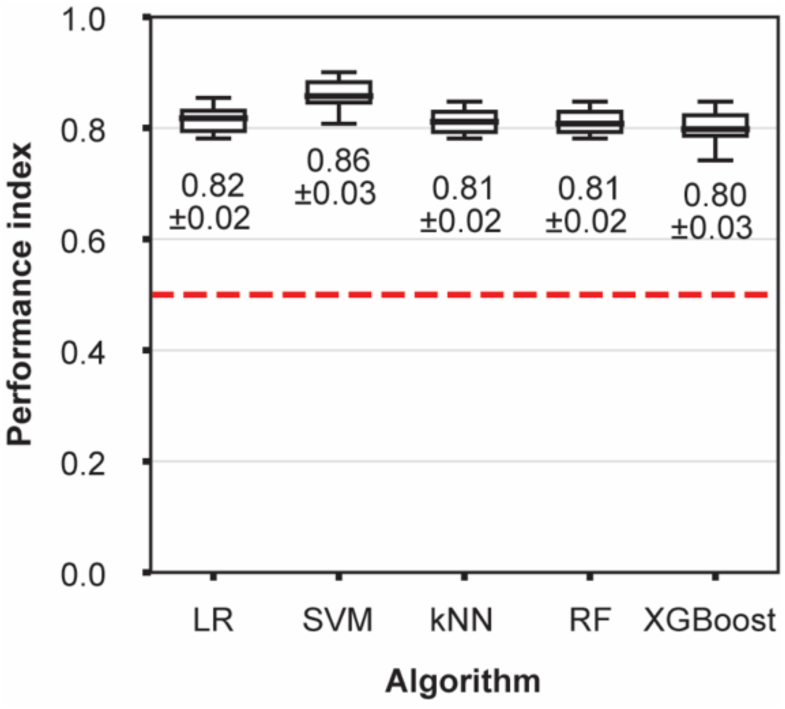
Evaluation of alternative classifiers of data presented in Figure 2G. LR=Logistic Regression, SVM=Support Vector Machines, kNN=k Nearest Neighbours, RF=Random Forest.

## Materials and Methods

### Fly husbandry

Flies were maintained on SYA medium (5% sucrose, Fisher Scientific BP220-10; 10% Brewer’s yeast (w/v), MP Biomedicals SR3010, lot no. S4707; 1.5% (w/v) agar, Sigma A7002; 0.3% nipagin (10% in EtOH), 0.3% propionic acid) in an insectary at constant 25oC, 12/12h light cycle. To set up experiments, eggs from an overnight egg lay were collected and suspended in PBS, and 20µl was pipetted onto 50ml SYA. After 10 days development, newly eclosed flies were transferred to fresh media. Flies were CO2 anaesthetised 48h later and sexed.

### Fly genetics

Where genotype is not specified, experiments were conducted with flies from the *Dahomey* background (gift from UCL Institute of Healthy Ageing) bearing the *w1118* mutation in the *white* gene locus.

*Drosophila* Genetic Reference Panel flies were purchased from the Bloomington *Drosophila* Stock Center, USA. The mitolines were described previously(*2*, *4*, *5*): briefly, flies from populations originating in Australia and Benin (formerly Dahomey) were used to generate mitolines, by crossing males with nDNA of interest to females with mtDNA of interest (n=45 of each sex), and iterating to purge ancestral nDNA and replace with focal nDNA. This procedure was used to generate all four combinations of mtDNA and nDNA, with each combination triplicated. At the time of experimentation the mitolines had been maintained in this way for hundreds of generations, which will have completed eliminated F0 nDNA.

### Diets

Diet experiments used the SYA formulation (as above). High fat diets were generated by adding 15% margarine (v/v). 5% and 20% yeast diets for DR experiments were generated by adding appropriate mass of yeast, with no further alterations to other ingredients, and making media up to final volume.

### ATR-FTIR

Two days post-eclosion, female flies were collected and allocated either to bottles containing SYA medium, or high-fat/DR media as above (n=50-100 flies unless stated otherwise). Flies were transferred to fresh food every 48-72 hours. At nine days post-eclosion, flies were flash-frozen on a weighing boat on dry ice for 1-2 minutes, before transfer to 15ml falcon tubes containing silica beads (separated by cotton), for two weeks’ desiccation. Samples were desiccated because water strongly absorbs infrared radiation, interfering with FTIR spectroscopy measurements.

After two weeks of drying in desiccation tubes, spectra were acquired in a dry-air purged Bruker Vertex 70 spectrometer (Bruker Corporation, Billerica, Massachusetts, USA) equipped with a Globar lamp, a Deuterated Lanthanum α Alanine doped Tri-Glycine Sulphate (DLaTGS) detector, a potassium bromide (KBr) beamsplitter, and a diamond Attenuated Total Reflectance (ATR) accessory (Bruker Platinum ATR Unit A225). The resolution of the instrument was 10 cm^−1^ and the sample and background measurements were averaged from 16 scans. Furthermore, the absorbance was measured from 400 cm^−1^ to 4000 cm^−1^. A new background measurement was obtained every 5 samples. Each desiccated fly was transported individually from the desiccation tube to the ATR crystal using forceps by its wings. Pressure was applied using the anvil of the ATR. Conditions processed rotated with every new tube of flies assayed, such that any environmental interference would be spread equally among conditions.

### Data analysis

The data files obtained by FTIR spectroscopy were imported and subjected to pre-processing by running a previously developed script *Bad Blood* (https://github.com/magonji/bad-blood)(*9*), discarding files with low-intensity peaks (spectrum intensity <0.1 at wavenumber 600 cm^−1^), abnormal background measurement (outliers at wavenumber 1900 cm^−1^), atmospheric interference (CO_2_ or water vapour-related interferences in region 3900-3500 cm^−1^).

ATR-FTIR spectral data were analysed in Python using the scikit-learn library. The pipeline comprised two stages: feature selection and classification. Informative spectral features were first selected using XGBoost. Selected features were then used to train five classifiers: logistic regression (LR), k-nearest neighbours (kNN), C-support vector machines (SVM), random forest (RF), and XGBoost. For each classifier, model parameters were tuned and the combination yielding the highest reclassification accuracy was selected.

Classifier performance was evaluated using k-fold cross-validation (k=20). In each iteration, data were split into a training set (80% of samples) and a validation set (20%). Stratified splitting ensured equal proportions of all conditions across training and validation sets, and data were shuffled at each split. This procedure generated distributions of correct and incorrect reclassification across 20 independent splits, enabling robust performance comparison among algorithms. For each classifier, the average accuracy across the 20 optimised models was computed and reported. SVM consistently outperformed alternatives across all analyses (see Supplementary Materials) and is reported in the main text unless otherwise stated.

To test the generalisability of the starvation-resistance classifier, the SVM trained on DGRP spectral data was applied to spectra from the independent mitoline panel (see Fly genetics). Each mitoline spectrum was assigned to the starvation-resistant or starvation-sensitive class based on its similarity to the corresponding DGRP-derived chemical fingerprints. Classification frequencies were computed per line and compared against observed starvation-resistance coefficients.

Data were plotted in Python using matplotlib and seaborn, or in R using ggplot2.

Phenotypic data were analysed in R 4.4.2. Survival data were analysed with a parametric survival model (rms::PSM). AIC values indicated that logistic distribution best fitted the survival data. Egg laying data were analysed with a negative binomial GLM (MASS::glm.nb). ANOVA tables were calculated with car::Anova, using Type-3 tests. Post-hoc tests were calculated with the EMMeans library. tSNE was analysed with Rtsne::Rtsne, with default parameters and random seed of 88.

